# Nightly dynamics of emotional content in dreams

**DOI:** 10.1101/2025.06.26.661683

**Authors:** Jessica Palmieri, Valentina Elce, Monika Schönauer

## Abstract

Emotional processing is a crucial adaptive function. Research suggests that sleep, particularly rapid eye movement (REM) sleep, may have a role in processing the emotional load of past events. Notably, dream experiences may offer insight into this nighttime process. Some studies have reported increased emotionality in dreams as the night progresses, possibly reflecting ongoing emotional processing in the sleeping brain. However, findings on how dream affect evolves throughout the night remain mixed. In this study, we investigated how emotional intensity in conscious experiences during sleep changes across the night and sleep stages. Participants (N_subjects_ = 20) were subjected to a multiple awakening paradigm, where they were awakened 4-5 times throughout the night and asked to recall their dreams (N_dreams_ = 61). Additionally, they rated the emotional intensity of their experiences using a structured cued questionnaire. Emotional intensity in dreams increased significantly throughout the night, with late-night dreams being more emotional than dreams collected during earlier sleep. Contrary to our expectation, this increase was not driven by dream reports obtained from REM sleep awakenings. Moreover, late-night dream reports were also significantly longer than those from early sleep, yet the length of the dream reports did not correlate with their emotional intensity. This suggests that the emotionality of dreams is not directly linked to the ability to recall the dream or its narrative complexity. Instead, it could be driven by emotional processes occurring independently throughout the night, or by other factors that regulate our access to dream experiences and their emotional content.

## 1. Introduction

*“In our dreams, images represent the sensations we believe they cause. We do not feel horror because we are threatened by a sphinx; we dream of a sphinx to explain the horror we feel.”* —J.L. Borges

As our brain transitions from wakefulness to drowsiness and finally to sleep, our conscious experience gradually shifts to an internally generated stream of images, sounds, and sensations. Grasping the underlying meaning of these nocturnal fabrications is often challenging. A number of theories attribute a function to dreaming, or regard it as a conscious manifestation of underlying adaptive processes, such as cognitive or affective processing (Crick & Mitchinson, 1983; Revonsuo, 2000; Valli & Revonsuo, 2009; Hobson, 2009; Hoel, 2021; Scarpelli et al., 2019). For instance, previous research suggests that dream construction may rely on memory (Baylor & Cavallero, 2001; Schredl & Hofmann, 2003; Nielsen & Stenstrom, 2005), and in other studies, dreaming has been linked to mechanisms of memory consolidation during sleep (Stickgold et al., 2001; Wamsley, 2014). While the relationship is unlikely to be linear or direct, growing evidence increasingly supports this connection (Wamsley et al., 2010; Wamsley & Stickgold, 2019; Schoch et al., 2019). Similarly, investigating dreams might provide a behavioral window to other cognitive or affective processes occurring during sleep.

### 1.1. Adaptive emotional processing during sleep

Sleep deprivation negatively affects emotional regulation during subsequent wakefulness: Sleep deprived participants showed higher impulsivity toward negative stimuli in a go/no-go task (Anderson & Platten, 2011) and increased negative emotional reactivity in response to disruptive (Zohar et al., 2005) as well as low-stress situations (Minkel et al., 2012). Emotional processing during sleep may thus be a crucial adaptive function (Goldstein & Walker, 2014).

Other studies have reported findings consistent with a regulatory emotional function of sleep, potentially reflected in dreaming. For instance, Sterpenich et al. (2019) found that individuals reporting higher levels of fear in dreams exhibited more efficient emotional regulation during wakefulness, a pattern reflected in stronger mPFC-amygdala connectivity. In a cross-cultural study, Samson and colleagues (2023) further investigated the adaptive value of emotionality in dreams. While emotionally charged dreams in non-clinical populations supported effective emotion regulation, this did not hold true for clinical groups, such as individuals with anxiety disorders. When fear extinction mechanisms are impaired, emotional content in dreams may thus fail to bring emotional resolution. In such cases, heightened emotionality in dreams might be indicative of a maladaptive emotional response. Taken together, this evidence suggests that dream affect could mirror ongoing emotion elaboration and regulation processes in the sleeping brain.

### 1.2. Dream characteristics and dream recall

Although dreaming occurs during all sleep stages (Rechtschaffen et al., 1963; Monroe et al., 1965; Brown & Cartwright, 1978; Crick & Mitchinson, 1983; Cavallero et al., 1992; Antrobus, 1995; Stickgold et al., 2001), dreams seem to exhibit different characteristics throughout the night. Dreaming from NREM sleep is reported as more thought-like and conceptual (W. D. Foulkes, 1966; Fosse et al., 2004), whereas REM dream reports usually appear more story-like, hallucinatory, and bizarre (Mamelak & Hobson, 1989; Hobson & Stickgold, 1994; Casagrande et al., 1996). Moreover, REM sleep is generally associated with longer reports (Antrobus, 1983; Cavallero et al., 1992; Hobson & Stickgold, 1994). The distinct neurophysiological characteristics of NREM and REM sleep may account for the differences in dream features between these stages (Diekelmann & Born, 2010). However, the literature remains divided on the nature of NREM and REM dreaming (Cicogna et al., 1991, 2000; Nielsen, 2000; Fosse et al., 2004; Oudiette et al., 2012), and not all theories agree on why dream reports from NREM and REM sleep may exhibit different features.

It is important to note that the formation of a dream memory follows the same phases as any other memory: after a dream is generated, it needs to be encoded, such that it can be successfully recalled (Nemeth, 2023). Some argue that the primary factor influencing dream recall lies in the distinct processes of dream generation that characterize NREM and REM sleep (Hobson et al., 2000), or in the varying intensity of dream generation throughout the night (D. Foulkes, 1999; Domhoff, 2018), which in turn would affect experience encoding and consequently, recallability (Fazekas et al., 2019). From these perspectives, dream reports may offer a relatively direct window into the complexity of the dream experience itself (Windt, 2016). Other views, however, suggest that the distinct mechanisms of memory processing in REM and NREM sleep (Diekelmann & Born, 2010) provide a neurophysiological environment that may be more or less conducive to the formation and retention of memory traces for the dream (Koulack & Goodenough, 1976; Goodenough, 1991). This would influence when and to what extent dream content can be retrieved, independent of the quality of the dream experience itself. These views likely complement each other: while dream generation may directly impact subsequent encoding, and thus, recallability, different cognitive mechanisms occurring during and after awakening from REM and NREM sleep may further affect the retrieval of dream content. The precise distinctions in dream characteristics across the night and sleep stages, as well as the factors underlying these differences, however, remain a subject of debate. One factor that might interact with sleep stages and shape nighttime dream dynamics is saliency. More bizarre, vivid, or emotionally salient dreams may facilitate encoding (Cipolli et al., 1993) and thus enhance subsequent recallability (Hartmann et al., 1991; Hartmann, 1996; Schredl & Montasser, 1996; Kunzendorf et al., 1997). This may also be promoted by bodily responses associated with dream intensity, as dream saliency appears to correlate with autonomic activity during sleep, such as eye movement density and breathing parameters (Nemeth, 2023).

Note that individual abilities to generate or accurately retrieve visuospatial imagery may also influence dream recall. Despite some contradictory findings (Waterman, 1991), dream recall has been associated with visuospatial skills and visual memory (Butler and Watson, 1985; Schredl et al., 1997). In a mirror tracing task (MTT), infrequent habitual dream recallers showed a lower baseline level of visuomotor skills compared to frequent recallers (Dumel et al. 2015). Taken together, these findings suggest that visuospatial skills may be crucial for either dream production or the ability to recall dream imagery.

Dream recall capacity has also been associated with fast sleep spindles, a well-established marker of memory consolidation during sleep (Cairney et al., 2018). Nielsen et al. 2017 found that fast spindle density during NREM stage 2 positively correlated with several measures of dream recall. Notably, spindle density was strongly associated with recall of negative dreams and the length of REM dream reports, as measured by word count. In line with these findings, Dumel et al. 2015 observed that overnight improvement in visuomotor skills was linked to negative dream affect in both frequent and infrequent dream recallers. Moreover, in a recent study, only participants who were able to recall dreams in the morning retained memory for negative pictures overnight —at the expense of neutral ones— whereas this advantage was absent in individuals who did not recall any dreams (Zhang et al., 2024). Collectively, these findings suggest a shared substrate possibly mediating dream recall, dream affect, adaptive processing of memories and adaptive emotional regulation.

### 1.3. Emotional processing and dream affect across the night

REM sleep seems to be particularly beneficial for emotional processing (Goldstein & Walker, 2014; Scarpelli et al., 2019; Wassing et al., 2019). Brain regions involved in emotional processing during wakefulness (e.g. hippocampus, amygdala, mPFC, anterior cingulate cortex) also characterize brain activity during REM sleep (Nofzinger, 2005; Dang-Vu et al., 2010; Nir & Tononi, 2010). The presence of REM sleep alterations in several affective and cognitive disorders (Tsuno et al., 2005; Nutt et al., 2008; Lamont et al., 2007; Palagini et al., 2013; Bersani et al., 2010; Wassing et al., 2019) further supports the view that REM sleep is crucial for efficient emotional processing and regulation. Importantly, this might be reflected in the content of dream reports collected from REM sleep awakenings. Previous studies suggest that REM dreams tend to be more emotional and perceptually vivid than NREM dreams (Foulkes et al., 1988; Merritt et al., 1994; Hobson et al., 2000; Kahn et al., 2002). However, a growing body of literature points toward a global increase of emotionality in dreams as the night progresses (Verdone, 1965; Sikka et al., 2014; Malinowski & Horton, 2021). Since the proportion of REM sleep increases across consecutive sleep cycles and is highest in the late night (Dement, 1957; Weber, 2017), this heightened emotionality may be due to either the increasing prevalence of REM sleep in the later sleep cycles or a more general rise in emotionality in dreams throughout the night, regardless of sleep stage.

Given the methodological constraints that may affect the assessment of emotionality in dream reports, and a number of contradictory findings regarding dream emotionality throughout the night (Verdone, 1965; Fosse et al., 2001; Malinowski & Horton, 2021; Sikka et al., 2014, 2017), further research is needed to clarify the progression of dream affect across the night and the role of different sleep stages in the emotionality of dreams.

### 1.4. Aims of this study

To investigate emotional processing during sleep from a behavioral perspective, we had participants sleep in the laboratory and repeatedly awoke them throughout the night to freely recall their dreams. At each of these awakenings, they had to answer questions regarding the emotionality of their dream experiences. This involved rating the intensity of positive and negative emotions present in their dreams on two separate scales. Emotionality was operationalized as the combined intensity of both positive and negative emotions experienced during dreaming. We then analysed how this measure evolved in the conscious experiences participants reported throughout the night, and whether dreams collected after REM sleep awakenings were rated more emotional than those obtained from NREM sleep awakenings.

Based on the predominant role attributed to REM sleep in emotional processing, we expected dream reports collected in the late-night, which is rich in REM sleep, to show higher emotional intensity than early-night dream reports. Moreover, we expected reports collected from REM awakenings throughout the whole night to carry more emotional content than dream reports collected from NREM awakenings.

REM dreaming has often been reported to be more story-like and associated with longer reports (Antrobus, 1983; Cavallero et al., 1992; Hobson & Stickgold, 1994). We thus also examined whether dream length varied across the night and different sleep stages, expecting overall longer dream reports in REM-rich sleep during the late-night, but also longer dream reports from REM awakenings compared to NREM awakenings across the whole night. Since emotionally salient dreams might be better encoded and thus recalled in more detail (Hartmann, 1996; Schredl & Montasser, 1996; Kunzendorf et al., 1997), we then tested the relationship between emotionality in dreams and the length of the reports.

## 2. Methods

### 2.1. Participants

This study included 20 participants (10 male) aged 20 to 30 years (25.5 ± 2.7 [mean ± SD]). All participants were healthy, non-smokers, and refrained from consuming alcohol, caffeine, or any medication—except for oral contraceptives—on the days of the experiment. They reported sleeping between 6 and 10 hours per night, maintained a regular circadian rhythm, and were neither extreme morning nor evening chronotypes, as assessed by the Munich Chronotype Questionnaire (MCTQ)(Roenneberg et al., 2007). None had engaged in shift work or undertaken long-distance travel in the six weeks prior to the experiment, and all were free of sleep-related disorders. Additionally, all participants were right-handed, as verified by the Edinburgh Handedness Questionnaire (Oldfield, 1971). Only individuals who frequently recalled dreams (at least three times per week, assessed by oral enquiry during the screening phase) were included in the study. The experiment received approval from the local ethics committee of the Department of Psychology at Ludwig-Maximilians-Universität München, and informed consent was obtained from all participants. One participant did not recall any dreams and was therefore excluded from all dream analyses (final sample: N = 19).

### 2.2. Experimental Procedure

This study was part of a larger project which investigated the incorporation of learning content into dreams. Please note that the current paper only reports those aspects of the study design and methodology that are relevant to the reported analyses. For further information on all methods, study rationale, and additional details on the dream reports, please see Kumral et al., 2023, Palmieri et al., 2025. Data for all dreams collected in this project is available in the DREAM database (Wong et al., 2023).

Each participant visited our laboratory twice: first for an adaptation night to familiarize themselves with the experimental procedure and environment, and then for the actual experiment. During the adaptation night, participants listened to an audiobook until they fell asleep. Simultaneously, a tone was played every 15 seconds, and participants were asked to press any key on a computer mouse to confirm they were still attentive. Once participants reached consolidated stage 2 sleep and stopped responding to the tone, the experimenter turned off the audiobook. The audio was played through external speakers at a volume of 45 dB ± 2.92 dB (mean ± SD), measured at the approximate location of the participant’s head in bed to ensure the story remained audible. On the experimental night, participants again fell asleep while listening to one of four randomly assigned audiobooks and were subjected to the same timed auditory stimulation of the adaptation night. The four audiobooks were selected from various genres and differed in themes, emotional engagement, and narrative style. However, the stories were not specifically chosen to influence participants’ emotionality. We presented the German versions of the commercially available audiobooks to participants, read by different male readers.

They were awakened after approximately one sleep cycle (87 ± 16 min [mean ± SD]) by the experimenter entering the room and addressing them by name. The time range was based on the approximate duration of a typical sleep cycle (∼90 minutes). However, both the audiobook manipulation and the multiple awakening paradigm may have affected sleep onset latency, increasing variability across participants. We relied on real-time EEG monitoring to ascertain that participants were asleep before initiating each awakening. If they did not wake up, the experimenter gently touched their arm while repeating their name. Participants were informed of this procedure in advance. Upon waking, they were asked to report their conscious experience just before awakening (see below for details on dream reports). They were then instructed to go back to sleep while resuming the audiobook from the point where they had left off. This process was repeated up to five times per night. The audiobook manipulation and keypress responses are not relevant to the results reported in this manuscript.

### 2.3. Dream reports

After each sleep period, participants were asked about their conscious experience before the moment of awakening. For this purpose, a standardized protocol was set prior to data collection and executed by instructions during each awakening. The overall procedure was an implementation of a commonly used method for dream enquiry (see Schoch et al., 2019). Dream reports were gathered systematically by repeatedly prompting participants—up to three times— to describe what had been on their mind just before waking. If a participant reported a conscious experience, they were instructed to provide a detailed account, specifying who was involved in the dream, where it took place, and what the action was. To encourage further recall, the experimenter asked, “Can you recall more?” up to three times. Following the free recall phase, participants completed a structured dream questionnaire designed to assess the characteristics of the reported dreams. This included predefined questions about the dream’s setting, characters, events, perspective, identifying the dream’s central theme, as well as emotional tone (see Table S8 for further information). To quantify emotional intensity, participants rated how positive and negative their dream felt on two separate scales ranging from 0 to 3 (see Table S9 and paragraph 2.5 for further information). All participants were required to respond to each question, but they could indicate if they had no memory of a particular aspect. The order of questions remained consistent across all trials to maintain a standardized procedure.

### 2.4. Polysomnography

Sleep EEG was recorded using a 128-channel active Ag/AgCl electrode system (ActiCap, Brain Products, Gilching, Germany) with a 1 kHz sampling rate and a high-pass filter set at 0.1 Hz. Electrode placement adhered to the extended international 10–20 system (Klem et al., 1999). For sleep staging, recordings were divided into 30-second epochs, and linear derivations were calculated (EEG: C3 and C4 referenced to the contralateral mastoids, as well as vertical and horizontal eye movements and EMG activity). Sleep stages were determined using electrodes C3 and C4 following standard criteria, with assessments conducted by two independent raters. Any discrepancies were resolved by a third rater.

### 2.5. Analysis

All statistical analyses were performed using RStudio 2021.09.1.

#### Emotionality in dreams

We explored the emotional tone of dream reports across the night. At each awakening, after freely describing their dream experience, participants were asked to rate how emotionally positive and emotionally negative each dream was perceived, using two standardized questions (Schredl & Doll, 1998). More specifically, to the question: “*Where there positive feelings in the dream?”,* they could answer with the options: “*No”, “Somewhat”, “Moderately”, “Strongly pronounced*”. The same question was repeated for negative emotions. The answers were later converted on a scale from 0 to 3. For each dream, these two separate scores were summed up to obtain an overall emotion score. In this study, emotionality in dreams was thus operationalized as the cumulative rated intensity of positive and negative emotions (*emotion score*).

To investigate the nightly dynamics of emotional content in dreams, we deliberately disregarded the valence of the reported emotions and focused primarily on their intensity. In subsequent analyses, we also examined the differential contribution of emotional intensity for positive and negative emotions, seperately. This was done to control for potential confounds in the emotional content of dreams that may have been introduced by our multiple awakening paradigm (such as a general deterioration of mood).

We initially examined whether emotionality scores differed significantly across audiobook conditions using a one-way ANOVA. Since no significant difference was found between the groups, data from the different audiobook conditions were pooled and treated as a homogeneous sample in all subsequent analyses (F(3, 48) = 1.88, p = 0.146; Eta2 = 0.10, 95% CI [0.00, 1.00], N = 52). The ratio of emotional versus non-emotional dreams was calculated for each awakening to assess the overall development of emotionality in dreams throughout the night (see Table S3). Only few participants woke up five times. Due to the low amount of overall dreams reported during the fifth awakening (N = 3), to avoid data misinterpretation it was not included in the descriptive assessments and visualizations of emotionality development across the night (fig. 1a, fig. S1) but it was included in all following analyses. A Wilcoxon signed rank test was then used to compare the *emotion score* in early and late awakenings (N = 10, two missing values for the *emotion score*). The first two awakenings of the experimental night were classified as early awakenings, while the last two were categorized as late awakenings. If a participant experienced five awakenings in total, the third awakening (i.e. the awakening in the middle of the night) was excluded from analyses comparing early and late awakenings. Following the same rationale, the difference in dream emotionality between NREM and REM awakenings was first calculated with a Wilcoxon-signed rank test. For this analysis, only participants who provided dream reports during both NREM (N2 and SWS) and REM awakenings were included. This resulted in a reduced dataset where dream report data were paired with the corresponding sleep stages for each participant (N = 7). Due to the reduced sample size after pairing the data, we also conducted exploratory analyses on the entire dataset to assess whether differences across night segments (N = 52; N_early_= 15, N_late_= 37, nine missing values for the *emotion score*), as well as between NREM and REM sleep (N = 41; N_NREM_ =30, N_REM_ =11, nine missing values for the *emotion score*), followed the same direction as those in the paired samples using Mann-Whitney U tests. A linear regression model (lm function) was then implemented to further investigate potential influences of sleep stages on the difference in emotionality levels found between dreams from early and late awakenings, selecting dreams reported after both NREM and REM awakenings (N = 41, 9 missing values). We selected the segment of the night and last sleep stage as predictors of the outcome variable emotion score (model used: *lm(emoscore_overall ∼night_segment*last_sleep_stage*). The variance inflation factor (VIF) analysis produced a score of 1.32 for the variable *night segment* and 3.41 for *last sleep stage*, suggesting no substantial collinearity issues that would warrant model exclusion. Given the low sample size, however, the results of the linear regression model should be interpreted with caution.

**Figure 1.**
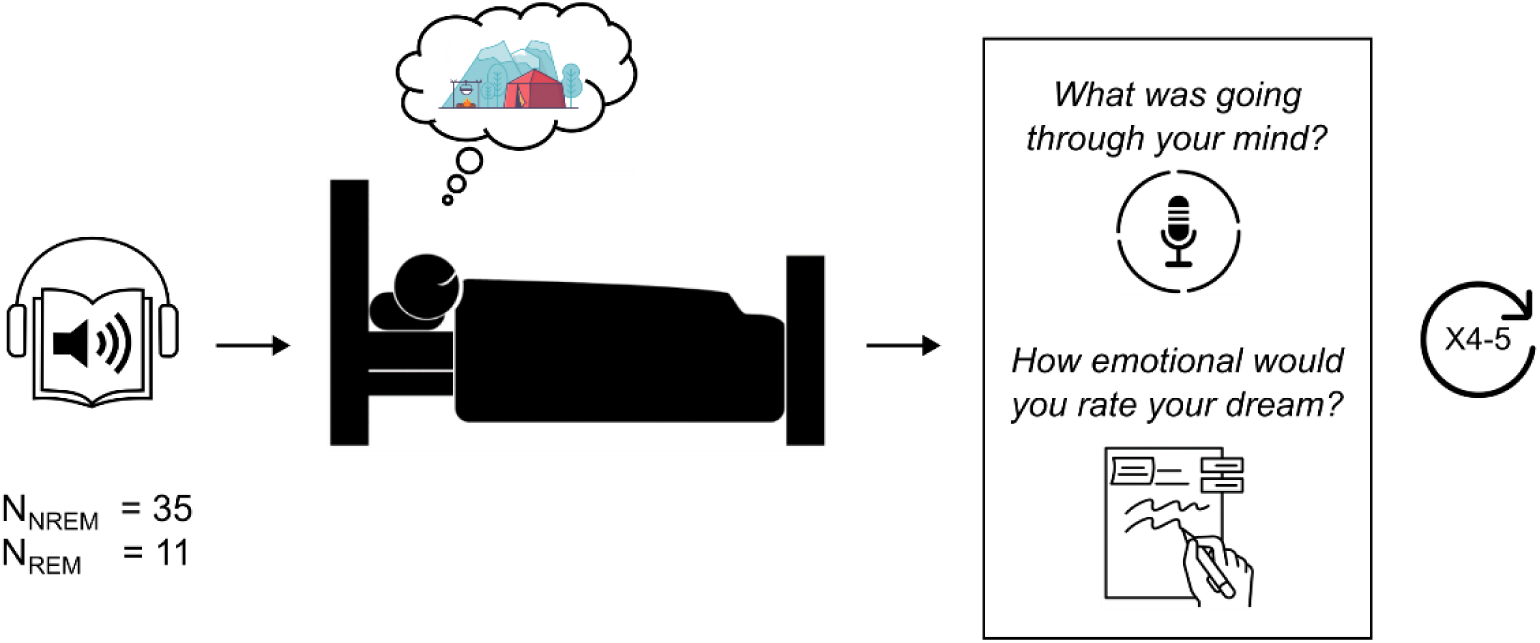
Experimental design. Participants listened to one of four audiobooks before going to sleep. Throughout the night, participants were awakened approximately every 90 minutes to describe their dreams and rate their emotional intensity. This procedure was repeated up to 4–5 times over the course of the night.

To partially control for a potential influence of the repeated awakenings on participants’ emotional state, we analyzed the development of positive and negative emotions in dreams across the night. For this purpose, we first compared positive and negative scores separately between early and late night and then examined whether there was any difference in emotional valence (positive vs. negative) during early and late night awakenings, using Wilcoxon-signed rank tests (N = 10, two missing values for the *emotion score*).

#### Dream length

Dream length was quantified as total number of words for each report. Dream reports were first redacted to include only content that was strictly inherent to the dream experience. Any commentary about the night or the task was removed.

Following the same approach used to assess group differences in emotionality scores, we employed the Wilcoxon signed-rank test to compare dream length across night segments (early vs. late sleep, N = 11, with one outlier exceeding the mean report length by 2 SDs excluded). As before, the first two awakenings were classified as early, while the last two were classified as late awakenings for each participant. Using Wilcoxon signed-rank tests, we then calculated the difference in dream length between NREM and REM awakenings, including only participants who reported dreams during both NREM and REM awakenings. This analysis was conducted on a reduced dataset with dream report data paired by sleep stage for each participant (N = 6, one outlier exceeding the mean report length by 2 SDs excluded). Similarly to the emotionality comparisons, pairing the data led to a reduced sample size. For this reason, we again conducted exploratory analyses using Mann-Whitney U tests on the entire dataset to assess differences across night segments (N = 56; N_early_= 18, N_late_= 38, three outliers exceeding the mean report length by 2 SDs excluded), and between NREM and REM sleep (N = 42; N_NREM_ = 33, N_REM_ = 9, three outliers exceeding the mean report length by 2 SDs excluded). To examine the potential influence of the last sleep stage on emotionality differences between dreams from early and late awakenings, we attempted to fit a linear regression model using dreams reported after both NREM and REM awakenings (N = 47). However, the model was rejected due to moderate to high multicollinearity between sleep stages and the interaction between *night segment* and *sleep stage* (VIF = 4.04 and 4.47, respectively).

In a secondary analysis, redacted dream reports were manually pruned to control for differences in narrative style across participants. In line with previous literature (Elce et al, 2025), this procedure included removing commentary about the dream experience itself, linguistic expressions used to introduce the experience as well as potential repetitions and self corrections. Following the same rationale as described above, we tested dream length differences across the night as well as its relation to dream emotionality based on the word count of these pruned reports.

#### Dream length and emotionality

We investigated whether dream emotionality was associated with the length of the corresponding dream report by correlating the overall *emotion score* and *dream length* for each report, using Kendall’s Tau (N = 47, with three outliers exceeding the mean report length by 2 SDs excluded and nine missing values for the *emotion score*).

We then applied a linear regression model (lm function) to assess whether emotionality could interact with the night segment factor to influence the length of dream reports. For this analysis, we used the absolute presence or absence of rated emotionality in dreams as emotionality variable. (N = 47, with three outliers and 9 missing values). *Night segment* and *emotionality* were selected as predictors of *dream length* (model: lm(*dream_words ∼ night_segment * emo_occurred*). Variance inflation factor (VIF) analysis produced a score of 2.1 for *night segment* and 3.2 for *emotionality*, suggesting no substantial collinearity concerns that would warrant excluding the model. However, due to the small sample size, the results of this linear regression should be interpreted with caution.

Finally, in a secondary analysis, we again used the dream length variable derived from the pruned version of the dream transcripts and performed a correlation between this measure of dream length and emotionality, using Kendall’s Tau (N = 48 with two outliers exceeding the mean report length by 2 SDs excluded and nine missing values for the *emotion score*).

#### Effect of sleep quality on dreams’ emotional content

To explore potential differences in participants’ sleep quality that may affect dream emotions, we used a median split approach and divided participants based on their average sleep efficiency across all awakenings. Since our multiple awakening protocol inevitably disrupts normal sleep architecture, this median split approach was preferred to the standard definitions of good or poor sleep efficiency. The median value for sleep efficiency in our pool of participants was 72%. For the analysis, participants with sleep efficiency values equal to 72 ± 1 (N = 3) were removed from the sample to reduce possible noise in the group comparison. We thus obtained a “higher sleep efficiency” group with values ranging from 79% to 93% (N = 8) and a “lower sleep efficiency” group with values ranging from 68% to 51% (N = 8). For each participant, the *emotion score,* as well as the individual *positive* and *negative scores* were averaged across all awakenings. We then used the Mann-Whitney U test to first compare overall emotionality (*emotion score*) between these two groups (N = 16). As it is been reported that “good” and “bad” sleepers differ on how intensely positive and negative emotions are re-expressed in dreams (Conte et al. 2021), we also compared *positive* and *negative emotion scores* within the two sleep efficiency group separately (for both groups, Npaired = 8), using the Wilcoxon signed rank test. Finally, a mixed-design ANOVA was conducted to assess the interaction between sleep efficiency group and emotion scores (positive vs. negative), with the sleep efficiency group as a between-subjects factor and the *positive* and *negative emotion scores* as a within-subjects factor.

## 3. Results

### 3.1. Descriptives

Each participant was awakened multiple times during the night (4.1 ± 0.44 [mean ± SD], N = 20). Across 82 awakenings, 61 dream reports were collected, resulting in a dream report rate of 74% (see Table S1-S2 for additional information). On average, participants reported 3.1 ± 1.7 [mean ± SD] dreams throughout the night, with some reporting more than one dream per awakening. Multiple dreams per awakening would entail different narratives, considered as distinct and temporally separated experiences by the participants. Likewise, multiple dreams would be associated with different answers in the structured dream questionnaire. One participant did not recall any dreams and was therefore excluded from all analyses (final sample: N = 19). During an average sleep period (across all awakenings), participants spent 7 ± 2 minutes awake, 8 ± 3 minutes in sleep stage 1 (S1), 29 ± 12 minutes in sleep stage 2 (S2), 14 ± 7 minutes in slow-wave sleep (SWS), and 16 ± 9 minutes in REM sleep. This resulted in an average total sleep time, between awakenings, of 67 ± 13 minutes [mean ± SD].

### 3.2. Dream emotionality

When completing their dream reports, participants were asked to rate the positive and negative emotional valence of each dream. Examples of dream reports emotionally negative, positive, and neutral can be found in Table S6. We summed these two scores to create an overall *emotion score* for each dream report. We first evaluated the development of emotionality in dreams throughout the night. The ratio of emotional dreams over non-emotional dreams more than doubled towards the end of the night (see Table S3, fig. S1-S2), indicating that later awakening dreams tended to be more emotional in nature compared to dreams from early sleep. This increase of emotionality in late night dreams was confirmed when comparing the emotion score between early and late sleep awakenings (Z = 4.5, p = 0.03*, CI[-2.08 -0.5], N_paired_ = 10, fig. 2b).

**Figure 2.**
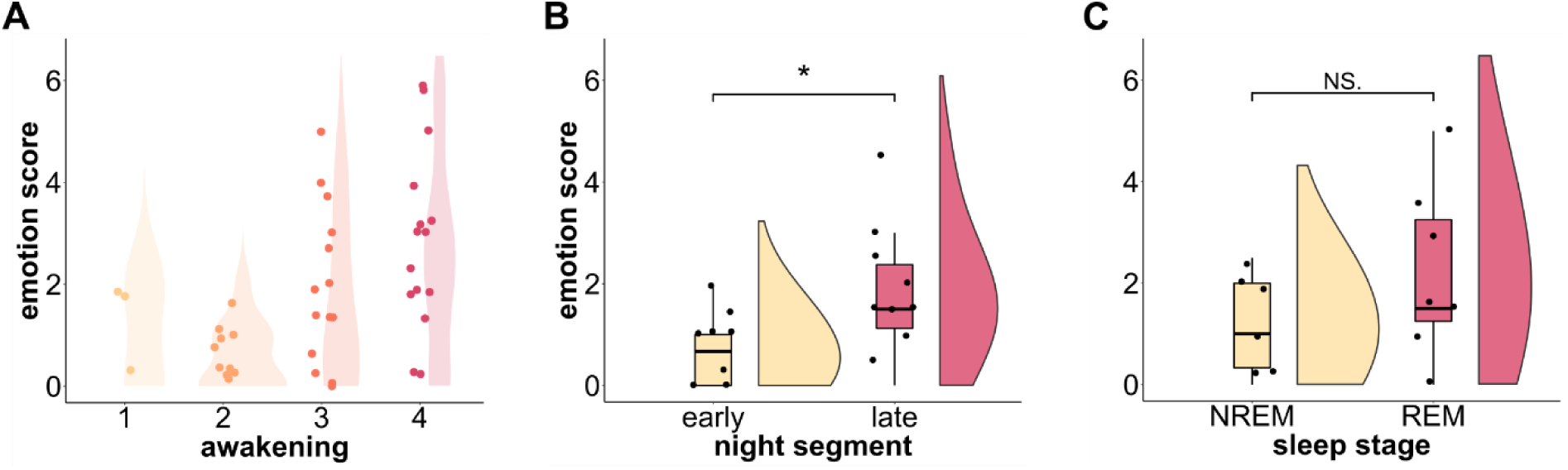
Overall emotionality in dreams across the night. A. Development of emotional dreams across the night. B. Comparison of the overall emotion score in dreams between early and late night. C. Comparison of the overall emotion score in dreams between NREM and REM awakenings. The difference is not significant (N.S). Due to the low sample size, data should be interpreted with caution. In the plots depicting group comparisons, the horizontal black lines within the boxplots represent the median value, whereas the dots represent the single observations. * p < 0.05.

Since the amount of REM sleep increases towards the end of the night (Dement, 1957; Weber, 2017) and REM sleep has been associated with emotional processing (Goldstein & Walker, 2014; Wassing et al., 2019), we compared the emotion score between dreams from REM and NREM awakenings. We could not observe any significant difference between the two groups (Z = 5, p = 0.25, CI [-3.83 1.25], N_paired_ = 7, fig. 2c). Due to the reduced sample size obtained when pairing the data by participant, results need to be interpreted with caution. To corroborate our analysis, we exploratively also ran group comparisons on the whole dataset. These analyses confirmed the results obtained from the paired data, showing a significant increase in emotionality scores in late sleep compared to early sleep (U = 156.5, p = 0.01*, CI[-2.00 0.00], N = 52 (N_early_= 15, N_late_= 37)), but no significant difference between NREM and REM awakenings (U=109.5, P=0.08, CI[-2.00 0.00], N = 41 (N_NREM_ =30, N_REM_ =11)).

Although the emotionality comparison across sleep stages was not statistically significant, reported emotionality was descriptively higher for dreams collected after REM sleep awakenings. For further investigation, we ran a regression model to test whether the occurrence of NREM or REM sleep before each awakening during early or late sleep could explain this difference in emotionality levels across the night. The model was statistically significant overall (R2 = 0.21, F(3, 37) = 3.37, p = 0.03*, adj. R2 = 0.15). However, none of the individual predictors or their interaction showed significant effects on emotionality (see Table S4 for further details). Note that there were overall fewer awakenings from REM sleep than from NREM sleep, which may have limited power for this analysis.

Furthermore, as the overall affect of participants might have deteriorated across the night because of the multiple awakenings, we also analyzed the positive and negative emotional load in dreams separately. A significant increase in negative emotions was found when comparing early and late awakenings, whereas this was not the case for positive emotions (negative: Z = 5, p = 0.04*, CI [- 2.00 0]; positive: Z = 6, p = 0.34, CI [-1.2 1.0], N_paired_ = 10, fig. 3a). However, negative emotions in late sleep dreams, as well as early sleep dreams, were not significantly higher than positive ones (late dreams: Z = 8, p = 0.12, CI[-1.50 0.5], N_paired_ = 10; early dreams: Z = 10, p = 0.5, N_paired_ = 10, fig. 3b).

**Figure 3.**
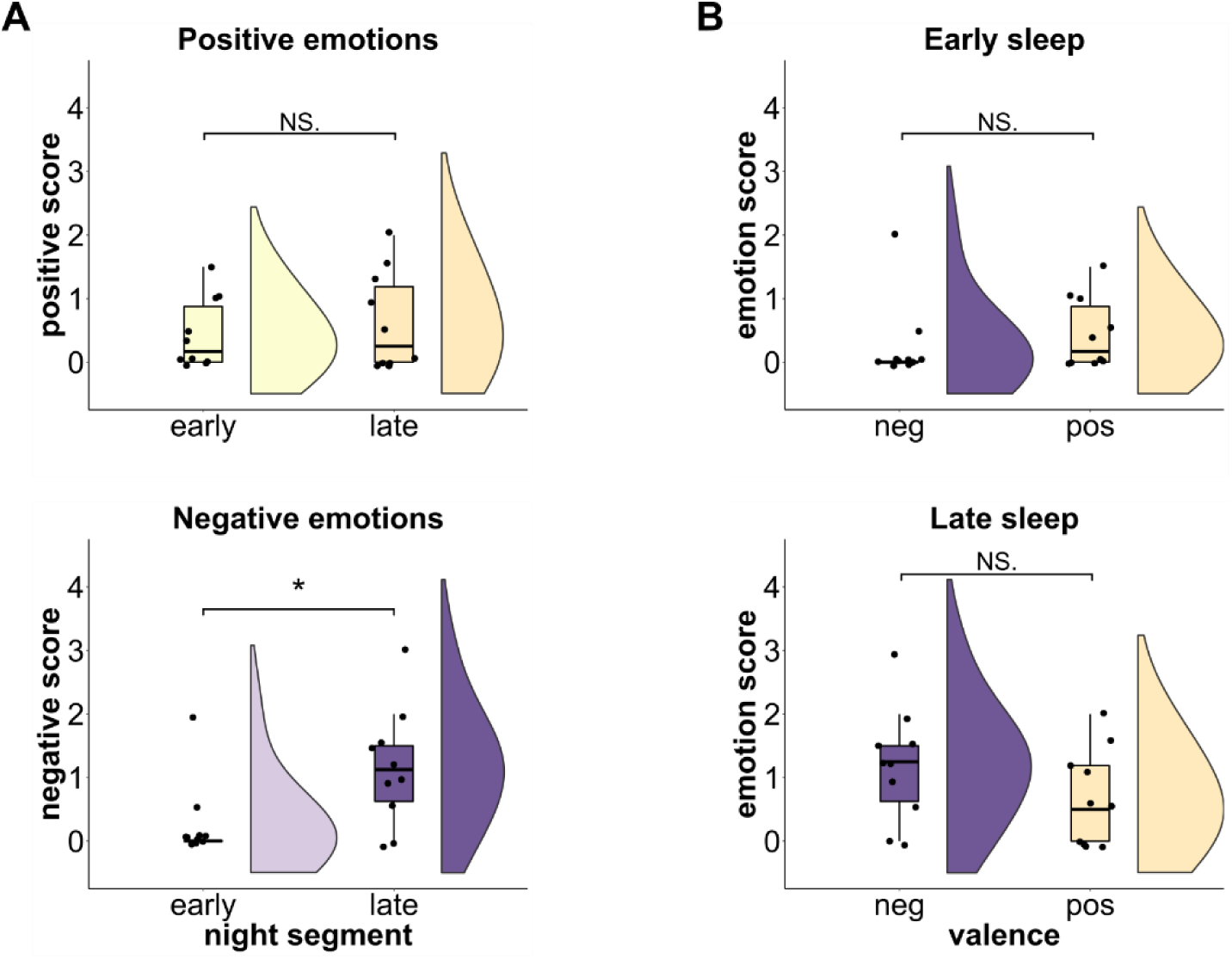
Positive and negative emotionality in dreams, night dynamics. A. Emotionality difference in dreams between early and late awakenings for positive scores (top left plot) and negative scores (bottom left plot). The difference for the positive scores is not significant (N.S). B. Difference between positive and negative emotionality in dreams during early sleep (top right plot) and late sleep (bottom right plot). Both differences are not significant (N.S). Due to the low sample size, data should be interpreted with caution. In the plots, the horizontal black lines within the boxplots represent the median value, whereas the dots represent the single observations. * p < 0.05.

### 3.3. Dream length

We quantified dream length as the number of words in each report. For all main analyses, we quantified dream length based on a redacted version of the dream reports, in which any commentary or reported content that was not strictly inherent to the dream experience itself had been removed.

Descriptively, dream length increased across awakenings (see Table S5, fig 4a, fig. S3). Following the same rationale adopted for assessing emotionality in dreams across the night, we first compared dream length between early and late sleep awakenings and found a significantly higher word count for dream reports collected in the late night compared to the early night (Z = 3, p = 0.008**, CI [-61.75 -16], N_paired_ = 11, fig. 4b).

**Figure 4.**
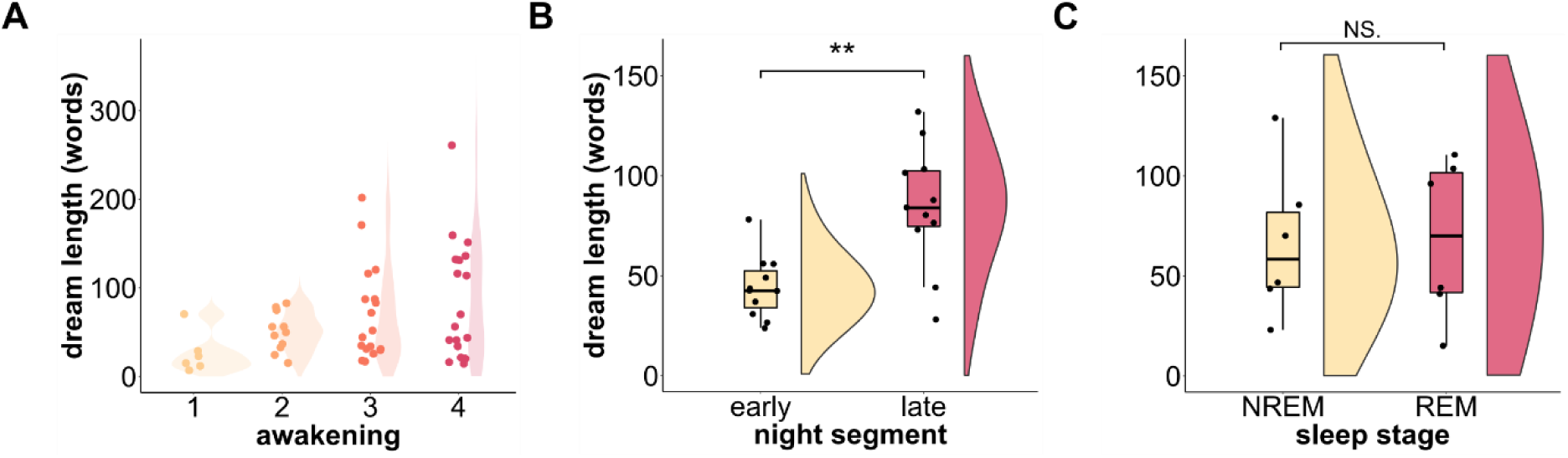
Length of dream reports across the night. A. Numbers of words in dream reports across awakenings. B. Comparison of the amount of words in dream reports between early and late night C. Comparison of the amount of words in dream reports between NREM and REM awakenings. The difference is not significant (N.S). Due to the low sample size, data should be interpreted with caution. In the plots depicting group comparisons, the horizontal black lines within the boxplots represent the median value, whereas the dots represent the single observations. ** p<0.01.

Dreams collected from REM sleep are usually more bizarre, story-like and elaborated than dreams collected from NREM (Mamelak & Hobson, 1989; Hobson & Stickgold, 1994; Casagrande et al., 1996). Since REM sleep is predominant in the late night, this could explain the length increase observed in late sleep. When testing the difference in dream length between reports collected from REM and NREM awakenings, however, we could not find any significant difference (Z = 9, p = 0.8, CI [-68.42 71.5], N_paired_ = 6, Table S6, fig. 4c). Also in this case, due to the reduced sample size following pairing data by participant, the results should be interpreted with caution. To strengthen our analysis, we again conducted exploratory group comparisons on the entire dataset.

These analyses confirmed the findings from the paired data, revealing significantly longer dream reports in late sleep compared to early sleep (U = 220.5, p = 0.03*, CI [-58.00, -2.00], N = 56; N_early_ = 18, N_late_ = 38). However, no significant difference was found between NREM and REM awakenings (U = 260, p = 0.38, CI [-35.00 14.00], N = 42; N_NREM_ = 33, N_REM_ = 9). Note that awakenings from REM sleep were fewer than those from NREM sleep overall, which may have reduced the statistical power of this analysis. Because of multicollinearity issues, we could not test whether waking up from REM or NREM sleep might explain longer dream reports in late sleep in a regression model (see Methods for further details).

Furthermore, to control for differences in narrative style that may have affected dream report length, we further pruned the dream reports (see Methods for further details) and derived a secondary dream length measure. Again, we performed group comparisons across night segments and sleep stages using this pruned dream length measure. Mirroring the main findings, dream length was significantly higher in the late night, with no significant differences emerging between NREM and REM sleep awakenings (see fig. S4 for further details).

### 3.4. Dream length and emotionality

We investigated whether the emotional intensity of each dream was associated with the length of the dream report by correlating the *emotion score* with *dream length*. The analysis, however, found no significant relationship (τ = 0.152, z = 1.38, p = 0.17, N = 47, fig. 5; see fig. S5 for further information). Although there appears to be no direct association between the rated emotional tone of dreams and dream length overall, the presence of an emotional component in dreaming may still interact with the night segment factor, contributing to longer dream reports in later sleep. To assess this, we fitted a linear regression model with *night segment* and *emotionality* as predictors of *dream length*. However, the model did not significantly explain the relationship. (R² = 0.11, F(3, 43) = 1.68, p = 0.185, adj. R² = 0.04, N = 47; see Table S7 for further details).

**Figure 5.**
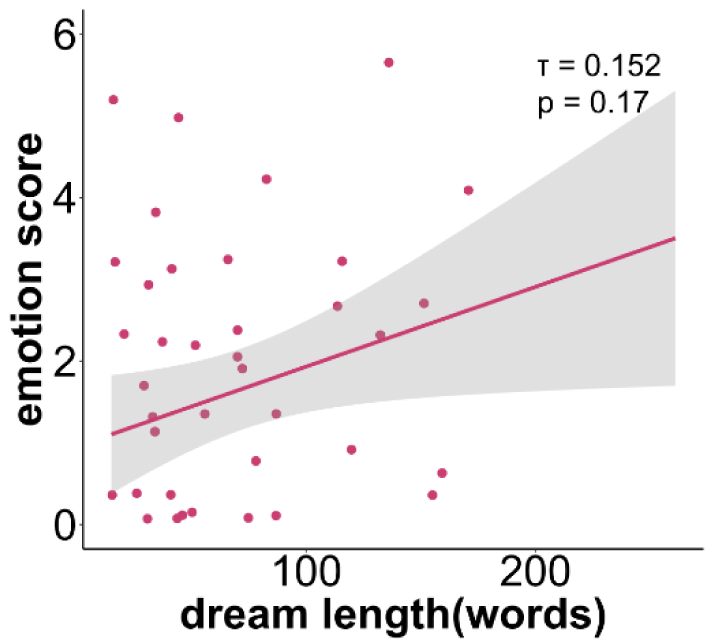
Relationship between emotionality and dream reports length. Kendall’s tau correlation between dream length and emotion score.

### 3.5. Sleep quality and emotionality in dreams

We ran an exploratory analysis to control for potential effects of sleep quality in the experimental night on dreams’ emotionality. For this purpose, participants were split into two groups based on their average sleep efficiency (see Methods and Table S2 for further details).

There was no significant difference in the overall *emotion score* between the two groups (U = 38, P = 0.5, CI[-1.19 2.0], N = 16; N_higher_ = 8, N_lower_ = 8, fig. S6). Within both groups, we next compared emotional intensity for *positive and negative scores*, as it is been shown that “good” and “bad” sleepers may display a different pattern of emotional re-expression in dreams for positive and negative emotions (Conte et al. 2021). However, we did not find any significant difference between *positive* and *negative* emotion scores for participants with lower or higher sleep quality during the experimental night (higher sleep efficiency group: Z = 38.5, P = 0.5, CI[-0.6 1.1], N_paired_ = 8; lower sleep efficiency group: Z = 33.5, P = 0.87, CI[-1.0 1], N_paired_ = 8; interaction effect between *sleep efficiency* group and *positive* and *negative scores*: F(1, 14) = 0.66, p = 0.431 N = 16).

## 4. Discussion

### 4.1. Emotionality in dreams increases in the late night

We examined emotionality in dreaming at different times during the night and across sleep stages. In our study, dream reports collected in the second part of the night were significantly more emotionally loaded than those collected in the early night. Despite some inconsistencies in the literature {Fosse et al., 2001}, this finding corroborates previous research suggesting that late-night dreams are more emotional than early-night dreams {Verdone 1965, Sikka et al. 2014, Malinowski & Horton, 2021}.

Literature agrees now that dreaming occurs across all sleep stages (Rechtschaffen et al., 1963; Monroe et al., 1965; Brown & Cartwright, 1978; Crick & Mitchinson, 1983; Cavallero et al., 1992; Antrobus, 1995; Stickgold et al., 2001), but its characteristics vary. Experiences reported from NREM sleep tend to be more thought-like and conceptual (Foulkes, 1966; Fosse et al., 2004), whereas REM dream reports are often longer, more hallucinatory, bizarre, and story-like (Mamelak & Hobson, 1989; Hobson & Stickgold, 1994; Casagrande et al., 1996).

Since REM sleep increases across sleep cycles (Dement, 1957; Weber, 2017), the observed increase in emotionality in the late night may correspond to the higher prevalence of REM sleep, which has been associated with more vivid and emotional content (Foulkes et al., 1988; Merritt et al., 1994; Hobson et al., 2000; Kahn et al., 2002). This explanation aligns with theories positing that REM sleep plays a critical role in emotional processing (Goldstein & Walker, 2014; Vandekerckhove & Cluydts, 2010; Perogamvros & Schwartz, 2015). Notably, brain regions involved in emotional processing during wakefulness, such as the amygdala, medial prefrontal cortex (mPFC), and anterior cingulate cortex, are also active during REM sleep (Nofzinger, 2005; Dang-Vu et al., 2010; Nir & Tononi, 2010). In some instances, these regions exhibit even higher metabolic activation than during wakefulness (Maquet et al., 1996; Braun et al., 1997; Nofzinger et al., 1997; Nir & Tononi, 2010). These dynamics may influence the emotional tone of dreams, amplifying their intensity during the late night.

Contrary to expectations, when comparing emotionality scores between NREM and REM awakenings, we did not observe any significant difference. Moreover, REM awakenings did not predict higher emotionality in late sleep in a regression analysis, thus suggesting that factors beyond REM sleep itself may contribute to the heightened emotionality of late-night dreams.

### 4.2. Dream length increases in the late night

We further found that dream reports were significantly longer in later sleep than in early sleep. Again, this may be due to the higher prevalence of REM sleep later in the night, as dream reports from REM sleep are typically longer (Antrobus, 1983; Cavallero et al., 1992; Hobson & Stickgold, 1994). Additionally, higher cortical activation during REM-rich late sleep (Dement, 1957; Antrobus et al., 1995) might enhance dream production, leading to dreams that are more perceptually vivid, bizarre, and story-like, ultimately resulting in longer reports (Mamelak & Hobson, 1989; Hobson & Stickgold, 1994; Casagrande et al., 1996). Note that longer dream reports might also indicate greater accessibility to the memory for the dream (Koulack & Goodenough, 1976; Goodenough, 1991), which could be state-dependent and vary across sleep stages.

When comparing dream report length between NREM and REM awakenings, however, we found no significant difference. REM awakenings may still have contributed to the increased dream report length in late sleep, as evidence suggests that especially late REM sleep leads to longer reports (Sikka et al., 2017). However, this could not be directly assessed in our dataset due to methodological limitations (multicollinearity between the variables, as well as small sample size, see Methods section 2.5, “*Dream length*” paragraph).

### 4.3. Dream length is not related with dream emotionality

In this study, we used freely recalled dream reports and subsequent emotional ratings of their content as an indicator of dreaming experience and its emotional characteristics. Dreams with strong emotional content, however, may lead to deeper encoding (Cipolli et al., 1993). As such, we explored the potential link between emotional intensity in dreams and the length of dream reports, hypothesizing that more emotional dreams could result in longer descriptions due to improved recallability (Hartmann et al., 1991; Hartmann, 1996; Schredl & Montasser, 1996; Kunzendorf et al., 1997). Longer reports for emotional dreams could also reflect richer verbalization not directly linked to the memory processing of the dream itself, but rather to the richness of speech used to describe emotional aspects of the dream (Sikka et al., 2014).

Contrary to expectations, we found no significant relationship between report length and rated emotionality. Despite the lack of direct association, emotionality might still have had an influence on the increased dream report length observed in the late night. To explore this, we tested whether the presence of emotional content in dreams could account for the increased length of late-night dream reports when included as a predictor in a linear regression model. However, the data did not support this hypothesis.

In conclusion, while shorter reports may provide less extensive descriptions of emotional features, this does not appear to correlate with how participants rated the perceived emotionality of their dreams. Conversely, more elaborate narratives did not necessarily convey stronger emotionality. Importantly, dream report length alone is not an accurate measure of the strength of the memory trace for a dream. Alternative and complementary indicators —such as participants’ confidence in the accuracy of their recall or the perceived ease of recall upon awakening— may better capture the accessibility and vividness of the memory trace. Recent theoretical accounts emphasize that subjective vividness or salience may reflect the precision of content-specific neural representations in the brain (Fazekas & Overgaard, 2016, 2018; Fazekas et al., 2020). According to this view, dream encoding and recall might be shaped by localized neural activity in brain regions specific to the dream’s content. Subject ratings, as a proxy for the precision of neural representations, may thus serve as additional, and potentially more sensitive indicators of dream memory strength compared to report length alone.

Overall, although both emotionality and dream length increased across the night, our findings suggest that these patterns do not directly mediate each other but may be shaped by distinct factors. It has been suggested that emotionality in dreams may generally increase throughout the night, independent of REM sleep (Verdone, 1965; Sikka et al., 2014; Malinowski & Horton, 2021). Some authors argue that this progression could be due to the accumulation of emotional processing occurring throughout sleep rather than being univocally dependent on specific sleep stages (Sikka et al., 2017). In line with this view, some studies showed that the characteristics of dream reports collected after NREM and REM sleep awakenings tend to become more similar towards the end of the night (see Malinowski & Horton, 2015).

### 4.4. Sleep quality and emotionality in dreams

It is worth considering that the sleep disruption introduced by our serial awakening protocol may have affected emotional processing and influenced the emotional tone of the dream experiences. While the use of such a paradigm was justified for the purpose of assessing emotional content in dreams throughout the night, it may also have disrupted emotional regulation as it would occur during an uninterrupted night of sleep. Moreover, prior research has identified a link between perceived sleep quality and the emotional content of dreams. For instance, individuals with insomnia or narcolepsy tend to report dreams with a more negative emotional tone compared to those of good sleepers (Feige et al., 2018; Pérusse et al., 2016; Fosse et al., 2002; Schredl et al., 1998;). Poor sleep quality has also been associated with a higher frequency of nightmares, suggesting that sleep fragmentation may increase negative emotionality in dreams (Levin et al., 1994), although the direction of causality remains unclear.

To partly account for the effect of the awakening protocol on participants’ emotional tone, we controlled for the progression of positive and negative emotions across the night. A greater intensity of negative emotions relative to positive ones as the task progressed could indicate that the repeated awakenings deteriorated participants’ mood. If this were the case, the overall increase in emotionality observed later in the night would be heavily confounded by the awakening protocol. Our analysis showed that the reported intensity of negative emotions was significantly higher in the late night compared to the early night. However, the rated intensity of negative emotions did not significantly differ from that of positive emotions, in either the early or late night. Although we cannot fully rule out potential biases introduced by the paradigm, this pattern of results does not suggest a substantial effect of the experimental protocol on participants’ mood progression. The observed increase in negative emotional intensity across the night may instead reflect emotional regulation processes, which may occur more prominently during REM-rich late sleep, or the late night per se. From an adaptive perspective, negative emotions —compared to positive ones—may benefit more from such processing (Conte et al. 2021).

It is important to highlight that individuals appear to exhibit different day and night profiles of emotional expression depending on their perceived sleep quality. Notably, Conte et al. 2021 found that good sleepers report more intense negative emotions in dreams than positive ones, potentially reflecting adaptive emotional processing. In contrast, poor sleepers do not show this difference, which may indicate an impairment in sleep-related emotional regulation. Conceptually, the findings by Conte et al. 2021 align with studies suggesting that emotional content in dreams may contribute to emotional resolution only when emotional processing mechanisms are intact (Sterpenich et al. 2019, Samson et al. 2023). Taken together, these findings underscore the role of sleep —and possibly of dreaming itself— in affective regulation processes.

In our study, we did not assess participants’ baseline perceived sleep quality. However, in an exploratory analysis, we divided the sample into two groups based on their sleep efficiency during the experimental night. The aim was to examine potential differences in overall dream emotionality, as well as expression of positive and negative emotions within the two groups as a function of sleep quality. We found no significant differences in overall emotionality between participants with higher versus lower sleep efficiency. Moreover, within each group, the intensity of positive and negative emotions did not significantly differ. Thus, in our study, sleep quality — operationalized as sleep efficiency during the experimental night — did not appear to influence dreams’ emotionality.

### 4.5. Limitations

It is important to acknowledge that the overall small sample size may have limited the statistical power of some analyses in this study. Specifically, more dream reports were collected after NREM sleep than after REM sleep awakenings. While this proportion mirrors the ratio of time spent in NREM and REM sleep during a regular night of sleep, it may have reduced the statistical power of analyses comparing effects of sleep stages on both dream length and dream emotionality. Considerably fewer measurements were available for REM sleep compared to NREM sleep, whereas the amount of data availabe for the early vs late night was more comparable. Note that the exact timing of dreaming before each awakening cannot be determined. Our analyses assume that dreams occurred within the few minutes before waking. However, our comparison of dream reports from NREM and REM awakenings does not rule out the possibility that emotional tone leaked into NREM or REM dreaming from preceding sleep stages.

Another important consideration is that the experiment was not designed to elicit a specific emotional state in participants. Although the audiobooks participants listened to while falling asleep may have varied in emotional engagement, this study did not systematically manipulate pre-sleep emotionality. Given that no effect of audiobook condition was found on overall dream emotional intensity, any inherent differences in emotional engagement across the narratives do not appear to have substantially influenced subsequent dream characteristics. This approach enabled us to quantify spontaneous emotional processes leaking in dreaming throughout the night. However, future research could address this limitation by manipulating the emotionality of pre-sleep learning material and examining how nocturnal dynamics of emotion processing in dreams relate to subsequent emotion regulation in wakefulness. Moreover, subjective ratings of dream emotionality may, in some cases, reflect emotions experienced in waking consciousness at the time of recall rather than those felt during the dream itself (Sikka et al., 2014). To more rigorously control for sleep-related trait-factors which may confound dream affect and emotional processing (Conte et al. 2021), perceived sleep quality should be considered as a potential confound in future studies.

In this study, we used the length of dream reports, measured by word count, as an indirect indicator of dream quality. While word count is commonly used as a proxy for dream length, it can be affected by individual differences in connected speech abilities, narrative style, verbal fluency, and the inclusion of content not directly related to the conscious experience during sleep. Although we redacted the dream reports to remove content not related to the dream experience, and applied a manual pruning procedure to remove further extraneous material in additional control analyses, this method remains a relatively coarse approximation. More fine grained linguistic approaches (Elce, Handjaras & Bernardi, 2021; Martin et al., 2020; Fogli, Aiello & Quercia, 2020; Cipolli et al., 2020; Casagrande, Cortini & Spoken, 2008) may provide a more nuanced and reliable assessment of dream structure and quality. Finally, we did not assess other relevant dream features such as bizarreness or narrative complexity, which could provide additional insights into dream quality.

## 5. Conclusions

We show that the emotional intensity of dreams increases across the night, irrespective of sleep stage, with late-night dreams being significantly more emotional than early-night dreams. This pattern may reflect ongoing emotional processing in the sleeping brain. Although the length of dream reports increased across the night, dream emotionality does not appear to be directly associated with the ability to recall the dream experience or its narrative complexity. Instead, it seems to be independently shaped by dynamic emotional processes unfolding throughout the night, or other factors regulating the access to our dream experiences and their emotionality.

## Acknoledgements

Funding: This work was supported by Deutsche Forschungsgemeinschaft (DFG, German Research Foundation) grants: SCHO1820/2-1, SCHO1820/5-1.

## Declaration of Competing Interest

The authors declare no conflict of interest.

## Authors Contribution

Conceptualization: JP, VE, MS

Methodology: JP, MS

Investigation: MS Visualization: JP

Funding acquisition: MS

Formal Analysis: JP, MS

Project administration: MS

Supervision: MS

Writing – original draft: JP, VE, MS

Writing – review & editing: JP, VE, MS

## Supplementary material

**Table S1.**
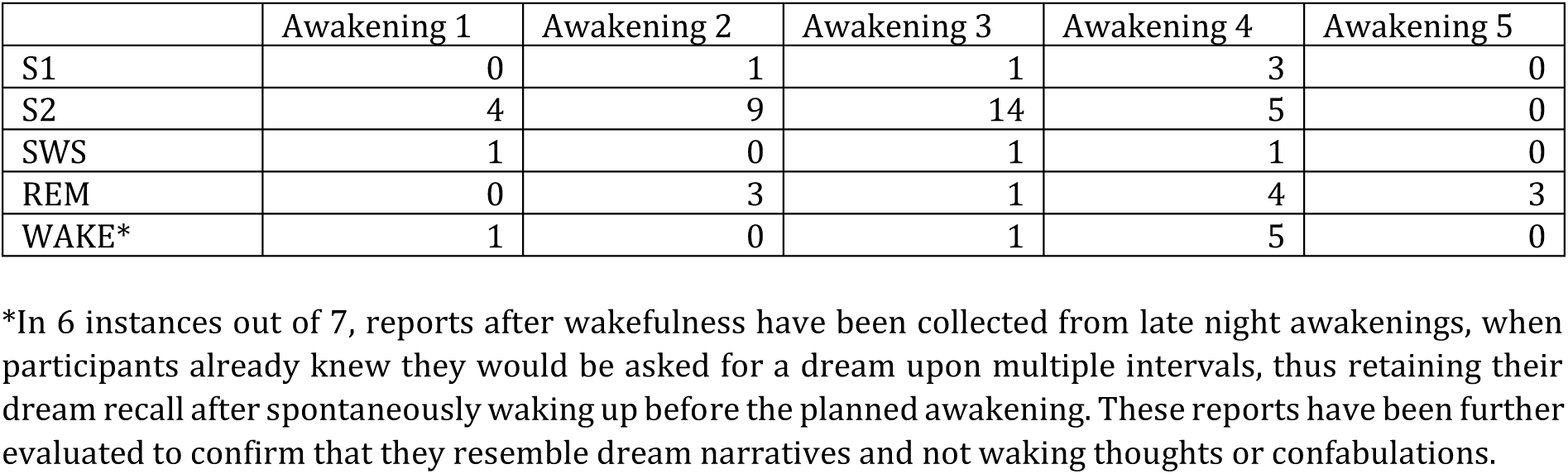
Distribution of last sleep stage by awakenings with dreams.

|  | Awakening 1 | Awakening 2 | Awakening 3 | Awakening 4 | Awakening 5 |
| --- | --- | --- | --- | --- | --- |
| S1 | 0 | 1 | 1 | 3 | 0 |
| S2 | 4 | 9 | 14 | 5 | 0 |
| SWS | 1 | 0 | 1 | 1 | 0 |
| REM | 0 | 3 | 1 | 4 | 3 |
| WAKE* | 1 | 0 | 1 | 5 | 0 |

\*In 6 instances out of 7, reports after wakefulness have been collected from late night awakenings, when participants already knew they would be asked for a dream upon multiple intervals, thus retaining their dream recall after spontaneously waking up before the planned awakening. These reports have been further evaluated to confirm that they resemble dream narratives and not waking thoughts or confabulations.

**Table S2.** Polysomnography data, by participant (averaged by awakening)

| Participant | Total sleep time in minutes<br>(mean ± SD, overall variance ) | WASO in minutes<br>(mean ± SD, overall variance ) | Sleep efficiency %<br>(mean ± SD, overall variance ) |
| --- | --- | --- | --- |
| 2 | 83 ± 20, 422 | 10 ± 11, 129 | 84 ± 15, 235 |
| 3 | 79 ± 10, 97 | 30 ± 24, 554 | 51 ± 13, 160 |
| 4 | 58 ± 29, 842 | 15 ± 8, 58 | 58 ± 20, 400 |
| 5 | 97 ± 30, 912 | 1 ± 1, 1 | 90 ± 9, 77 |
| 6 | 60 ± 43, 1867 | 11 ± 15, 222 | 65 ± 34, 1168 |
| 7 | 59 ± 41, 1682 | 20 ± 18, 307 | 55 ± 24, 596 |
| 8 | 75 ± 32, 1052 | 3 ± 3, 9 | 73 ± 16, 261 |
| 9 | 65 ± 68, 4610 | 13 ± 13, 169 | 53 ± 33, 1076 |
| 10 | 82 ± 37, 1385 | 2 ± 3, 7 | 93 ± 3, 11 |
| 11 | 70 ± 43, 1833 | 2 ± 1, 1 | 82 ± 22, 484 |
| 12 | 70 ± 31, 969 | 17 ± 20, 401 | 67 ± 19, 347 |
| 13 | 106 ± 40, 1618 | - | 93 ± 4, 13 |
| 14 | 76 ± 43, 1822 | 1 ± 2, 3 | 73 ± 13, 170 |
| 15 | 41 ± 36, 1321 | 2 ± 2, 4 | 50 ± 8, 57 |
| 16 | 70 ± 40, 1618 | 1 ± 1, 1 | 68 ± 26, 702 |
| 17 | 80 ± 29, 834 | 9 ± 17, 276 | 80 ± 12, 148 |
| 18 | 96 ± 30, 922 | 8 ± 17, 281 | 86 ± 11, 121 |
| 19 | 57 ± 28, 762 | 5 ± 8, 66 | 79 ± 13, 174 |
| 20 | 52 ± 32, 1044 | 6 ± 5, 26 | 63 ± 24, 597 |
| 21 | 51 ± 18, 338 | 3 ± 4, 16 | 71 ± 15, 224 |

**Table S3.** Emotionality in dreams across awakenings.

|  | Awakening 1 | Awakening 2 | Awakening 3 | Awakening 4 | Awakening 5 | All |
| --- | --- | --- | --- | --- | --- | --- |
| Neutral dreams | 2 | 6 | 6 | 4 | 1 | 19 |
| Emotional dreams | 2 | 5 | 11 | 13 | 2 | 33 |
| Positive emotions | 2 | 4 | 6 | 8 | 1 | 21 |
| Negative emotions | 1 | 1 | 9 | 11 | 2 | 24 |
| Emo vs non emo dreams (ratio) | 1.0 | 0.8 | 1.8 | 3.25 | *- | 1.7 |

\* Only three dream reports were collected from the fifth awakening. Thus, it was excluded from the descriptive assessment of emotionality development across the night.

**Table S4.** Effect of sleep stage and time of the night on emotionality in dreams, linear regression.

| Predictor | $\beta$<br>(Estimate) | t(37) | p-value | 95% CI |
| --- | --- | --- | --- | --- |
| Intercept (early sleep, NREM sleep) | 0.73 | 1.65 | 0.11 | [-0.17 1.62] |
| Late sleep | 0.48 | 0.87 | 0.39 | [-0.64 1.61] |
| REM sleep awakenings | -0.06 | -0.06 | 0.95 | [-2.00 1.87] |
| Late sleep*REM sleep awakenings | 1.60 | 1.41 | 0.17 | [-0.70 3.90] |

Summary: R<sup>2</sup> = 0.21, F(3, 37) = 3.37, p = 0.03\*, adj. R<sup>2</sup> = 0.15

**Table S5.** Average dream length across awakenings (outliers excluded)

|  | Awakening 1 | Awakening 2 | Awakening 3 | Awakening 4 |
| --- | --- | --- | --- | --- |
| Average words in dreams | 26 | 51 | 68 | 86 |
| SD (±) | 23 | 21 | 53 | 67 |
| Average words in dreams (pruned) | 21 | 44 | 53 | 65 |
| SD (±) | 25 | 17 | 47 | 53 |

**Table S6.**
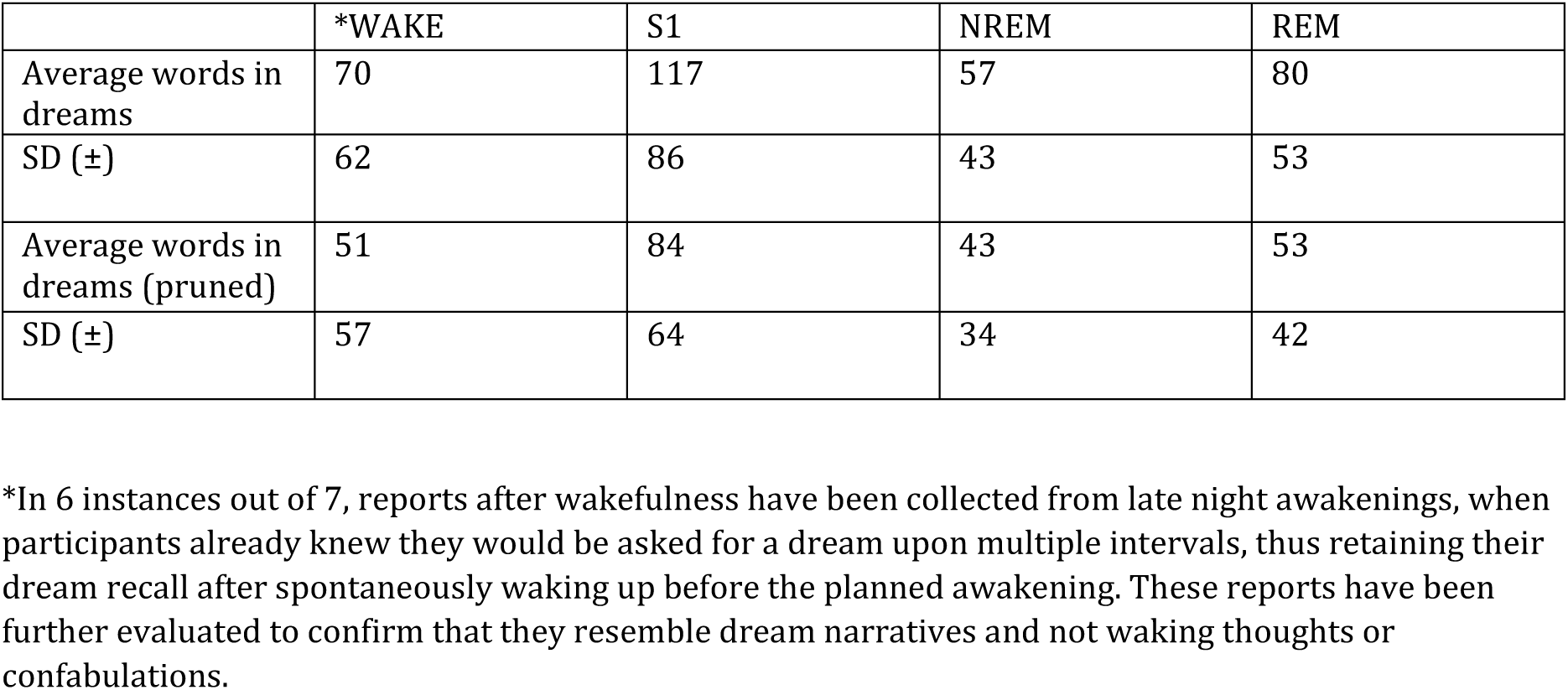
Average dream length by last sleep stage (outliers excluded)

**Table S7.** Effect of time of the night and emotionality on length of dream reports, linear regression.

| **Predictor** | **$\beta$**<br>**(Estimate)** | **t(43)** | **p-value** | **95% CI** |
| --- | --- | --- | --- | --- |
| Intercept (early sleep, neutral dreams) | 45.37 | 2.39 | 0.021* | [7.15, 83.60] |
| Late sleep | 24.33 | 0.96 | 0.34 | [-26.96, 75.61] |
| Emotional dreams | 5.13 | 0.18 | 0.86 | [-53.27, 63.52] |
| Late sleep*emotional dreams | 13.35 | 0.38 | 0.71 | [-57.97, 84.67] |

Summary:
$R^2 = 0.11$ , $F(3, 43) = 1.68$ , $p = 0.185$ , adj. $R^2 = 0.04$

**Table S8.** List of questions from the structured dream questionnaire.

| **Question** | **Answering options** |
| --- | --- |
| **PEOPLE-RELATED** |  |
| *Were there people in your dream? How many?* | *No - yes (state how many)- I don't know* |
| *Did you recognize the people?* | *No - yes - I don't know* |
| *Who were the people? Can you tell me more about them? (optional if people were in the dream)* | *Open answer* |
| *What did the people do? Can you tell me more about them? (optional if people were in the dream)* | *Open answer* |
| **ENVIRONMENT-RELATED** |  |
| *We will now ask you about the environment in your dream. Were you inside a building or outside?* | *Inside - outside* |
| *Were you in a room? (if inside)* | *No - yes- I don't know* |
| *Were you in a street? (if outside)* | *No - yes- I don't know* |
| *Were you out in nature? (if outside)* | *No - yes- I don't know* |
| *Were there buildings? (if outside)* | *No - yes- I don't know* |
| *Were there multiple buildings? If yes, how many? (if outside)* | *No - yes (state how many)- I don't know* |
| *Were you in a city? (if outside)* | *No - yes- I don't know* |
| *Were there any objects there? If so, how many and which ones? (for both inside and outside)* | *No - yes (state how many and which ones)- I don't know* |
| *Please describe the environment in more detail* | *Open answer* |
| *Did you recognize the environment?* | *No - yes- I don't know* |
| **DREAM PLOT-RELATED** |  |
| *Did you take part in the dream story yourself?* | *No - yes- I don't know* |
| *Which people/animals took part in the action? What did they do?* | *Open answer* |
| *Which objects were part of the action? What was done with them?* | *Open answer* |
| *What steps were taken?* | *Open answer* |
| *What was in the center of the field of vision?* | *Open answer* |
| *From which angle did you see the action (wide angle/zoom)?* | *Open answer* |
| **EMOTIONS-RELATED** |  |
| *Were there any positive feelings in the dream?* | *No - somewhat - moderately - strongly pronounced* |
| *Were there any negative feelings in the dream?* | *No - somewhat - moderately - strongly pronounced* |
| **CENTRAL ELEMENT** |  |
| *What would you describe as the central element of your dream?* | *Open answer* |
| **TIME OF DREAMING** |  |
| *Did you dream that right before I woke you up?* | *No - yes- I don't know* |
| *Did we wake you up from the dream?* | *No - yes- I don't know* |
| *Or was the dream already complete when we woke you up?* | *No - yes- I don't know* |
| *Can you tell us when you dreamt this?* | *Open answer* |

Note: This questionnaire was provided after participants freely recalled their dream report. Questions are listed in the original order of presentation.

**Table S9.** Examples of dream with and without emotional component.

|  | Translated version |
| --- | --- |
| Rated as negative dream | *"I dreamt of the war, there was a trench, I had a rifle and wanted to shoot but I couldn't; the dream was very emotional and negative, I didn't know the landscape and the people there but there were other people."* |
| Rated as positive dream | *"I was playing tennis with someone I knew from before, but I haven't talked to him in 15 years; it was next to a huge football pitch, there was an important match; we walked past after our tennis match and it was raining heavily and people were cheering: It was like a stadium, but there was no actual stadium; we then walked around the whole pitch and that's when I saw a mate of mine who I actually knew from tennis and he had accepted the ball and played it back into the pitch, once he appeared."* |
| Rated as neutral dream | *"I'm in a bed camp that was somehow on a school trip and I'm one of many children and one of them is reading aloud; and we have to stay there until he stops, but he never stops; and everyone else is already packing up, but I'm like: no, the story isn't over yet; that was inside."* |

Note: In this table, we report only a selection of dream reports that were rated as exclusively positive, exclusively negative, or neutral, for illustrative purposes. However, among all dream reports containing emotional content, some were rated ambivalently—that is, as containing both positive and negative emotions, either to the same or to differing degrees of intensity.

**Figure S1.**
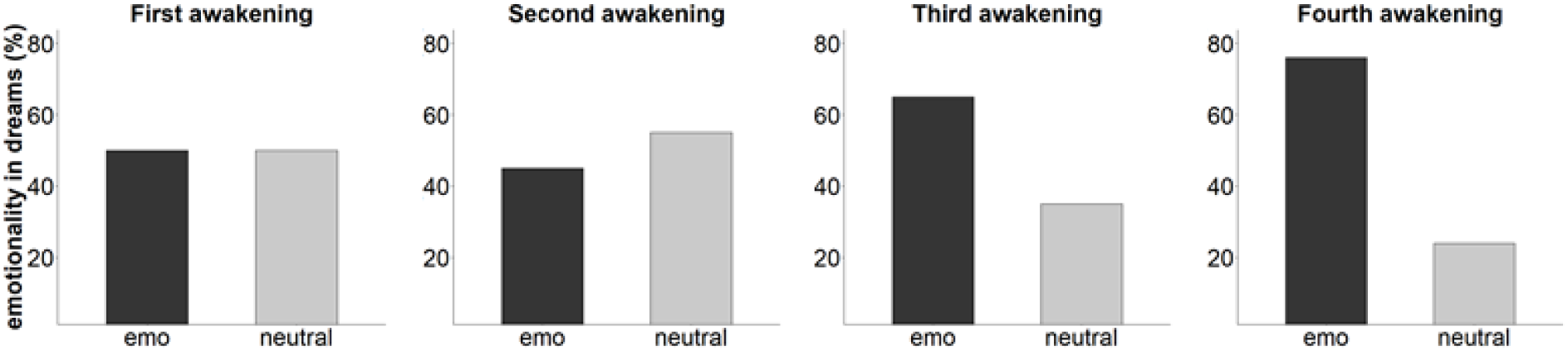
Proportion of emotional and neutral dreams during first to fourth awakening, in percent. Note: Only three dream reports were collected from the fifth awakening. Thus, it was excluded from the descriptive assessment of emotionality development across the night.

**Figure S2.**
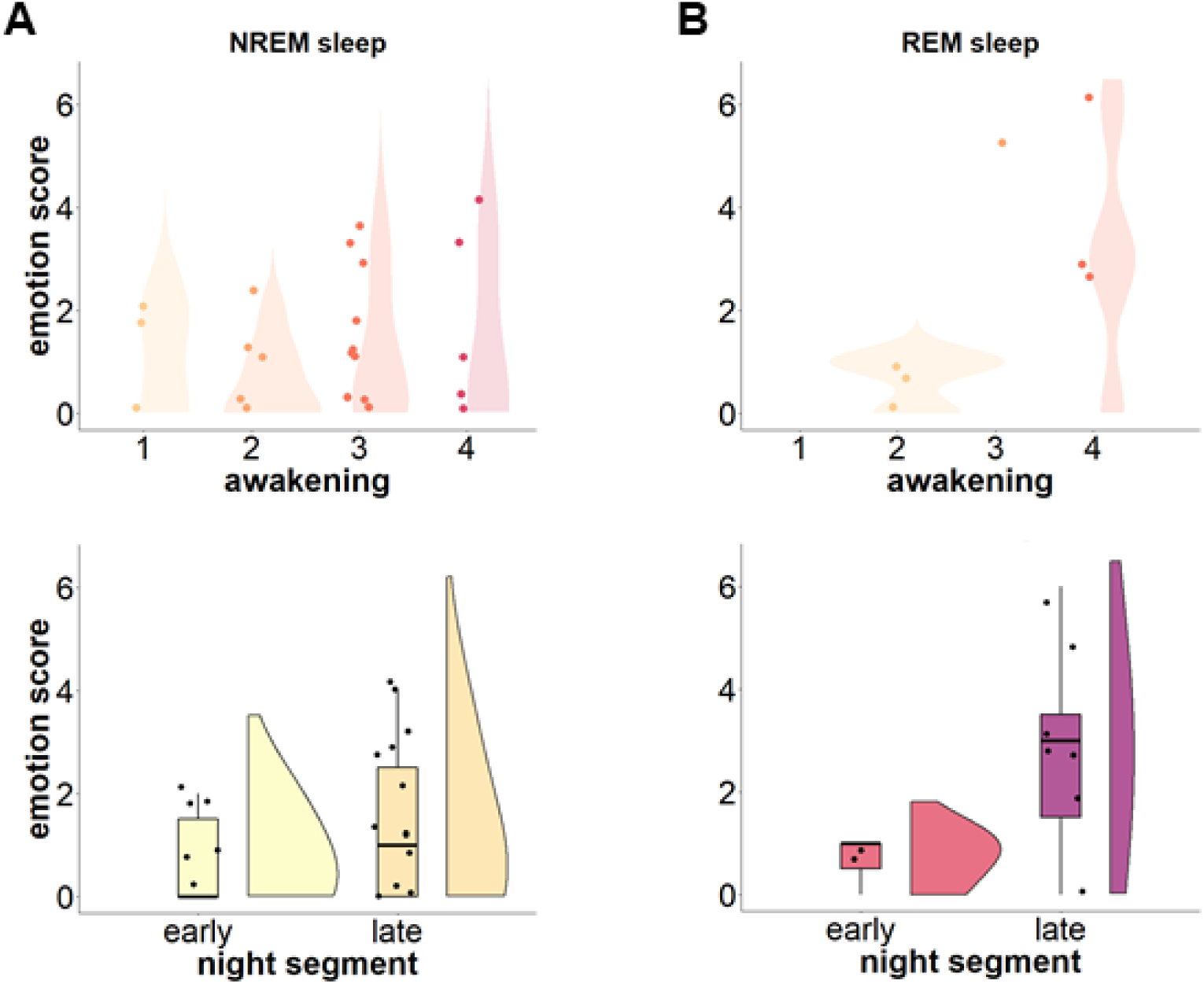
Emotionality in dreams, nightly dynamics by sleep stage. A. Emotion score in NREM sleep across awakenings (top left plot) and night segments (bottom left plot) over the whole sample. B. Emotion score in REM sleep across awakenings (top right plot) and night segments (bottom left plot) over the whole sample. In the boxplots, the horizontal black lines within the boxplots represent the median value, whereas the dots represent the single observations. No statistical tests were performed, given the very low sample size that would have resulted from pairing the data.

**Figure S3.**
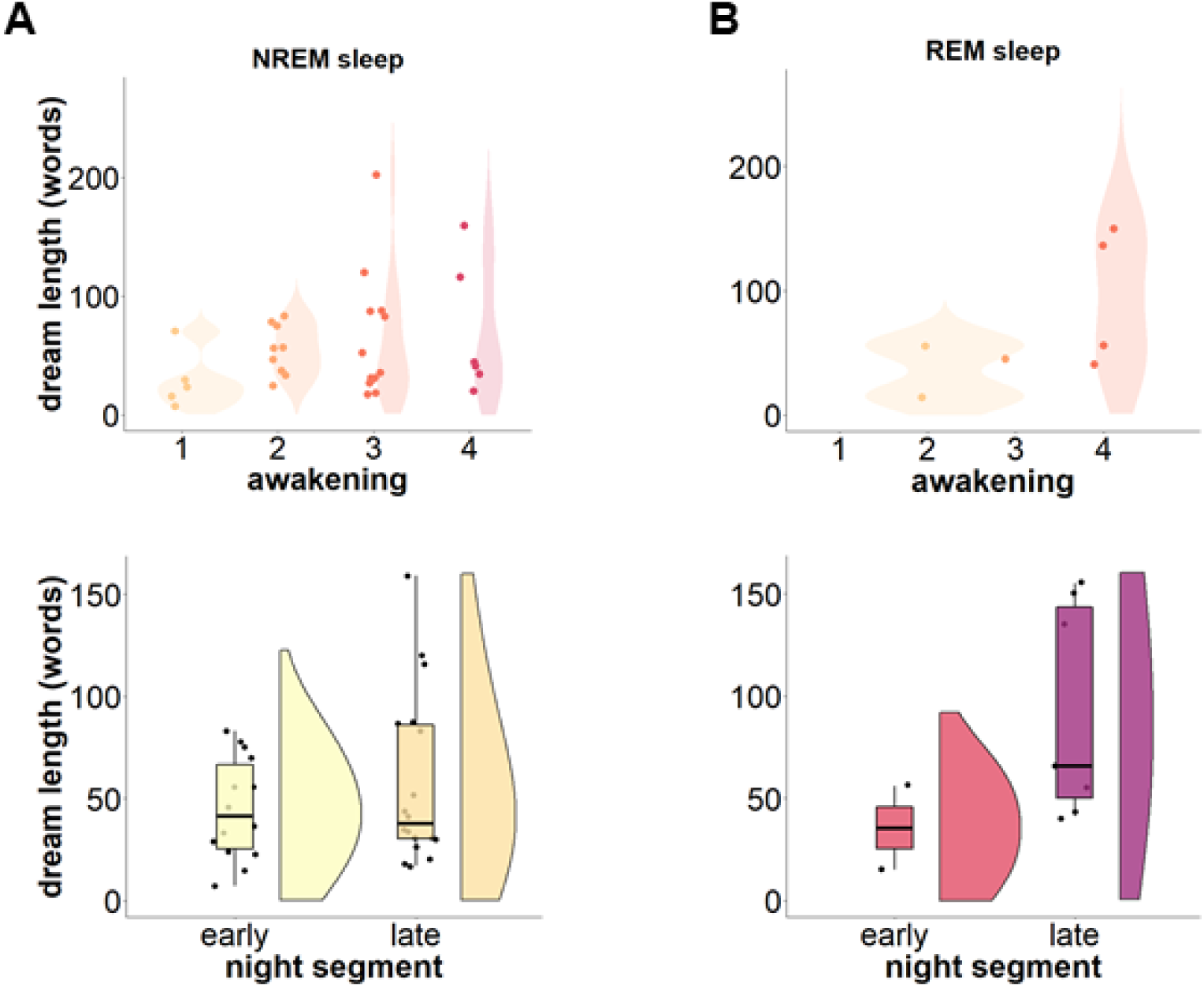
Dream length, nightly dynamics by sleep stage. A. Dream length in NREM sleep across awakenings (top left plot) and night segments (bottom left plot) over the whole sample. B. Dream length in REM sleep across awakenings (top right plot) and night segments (bottom left plot) over the whole sample. In the boxplots, the horizontal black lines within the boxplots represent the median value, whereas the dots represent the single observations. No statistical tests were performed, given the very low sample size that would have resulted from pairing the data.

**Figure S4.**
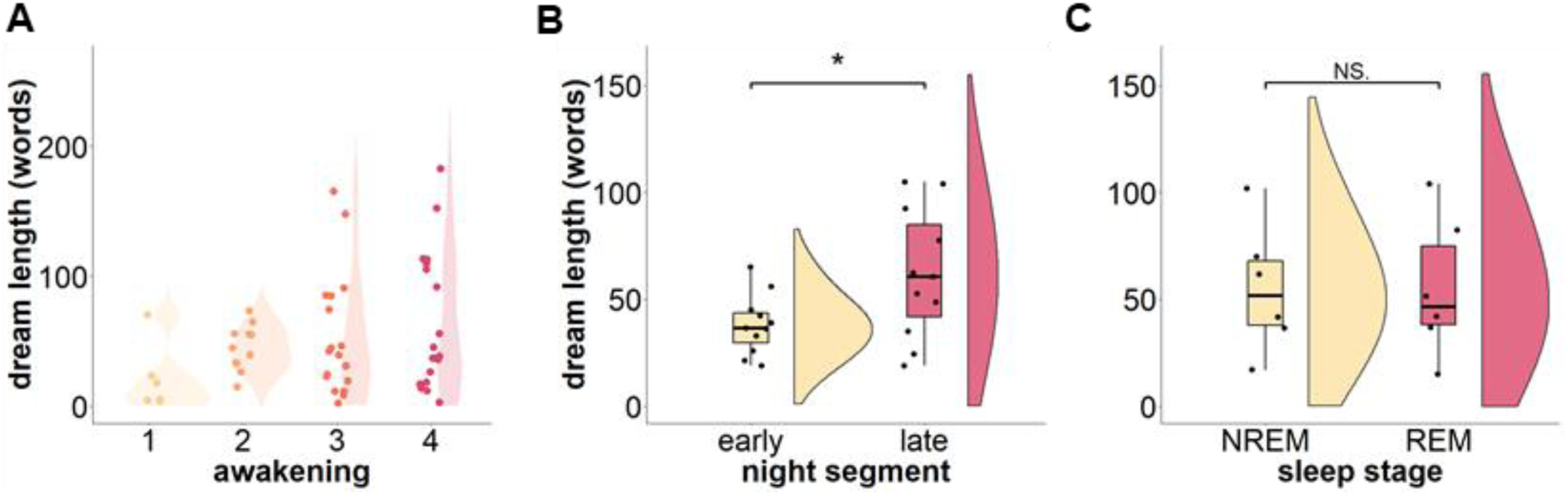
Length of dream reports (pruned version) across the night. A. Numbers of words in dream reports across awakenings. B. Comparison of the amount of words in dream reports between early and late night (Z = 9, P = 0.04*, CI [-42.0 -1.9], Npaired = 11). C. Comparison of the amount of words in dream reports between NREM and REM awakenings (Z=9, P=0.8, CI [-66.42 60], Npaired = 6). The difference is not significant (N.S). Due to the low sample size, data should be interpreted with caution. In the plots depicting group comparisons, the horizontal black lines within the boxplots represent the median value, whereas the dots represent the single observations. * p<0.05.

**Figure S5.**
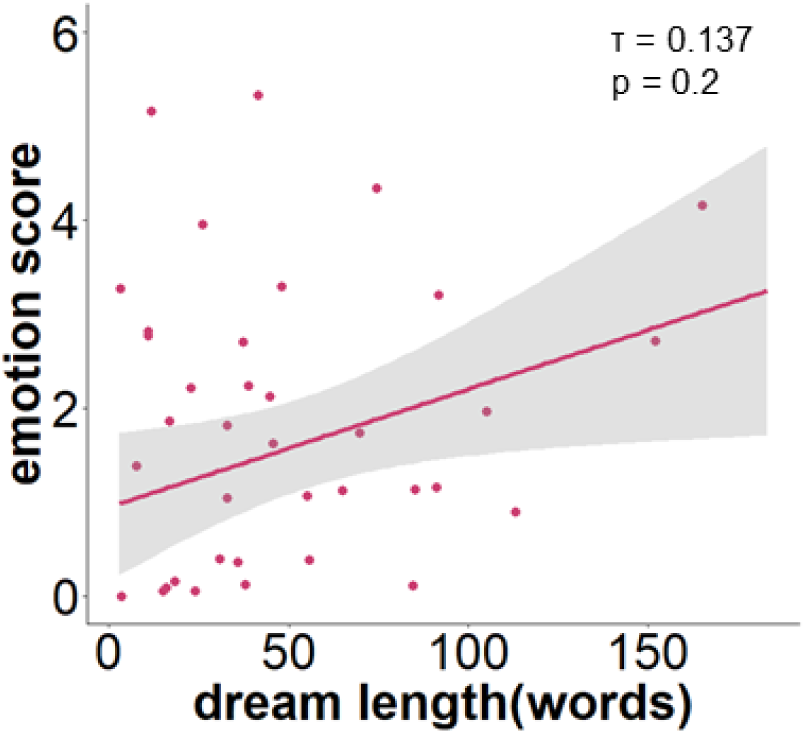
Relationship between emotionality and dream reports length (pruned version) Kendall’s tau correlation between dream length and emotion score (N = 48).

**Figure S6.**
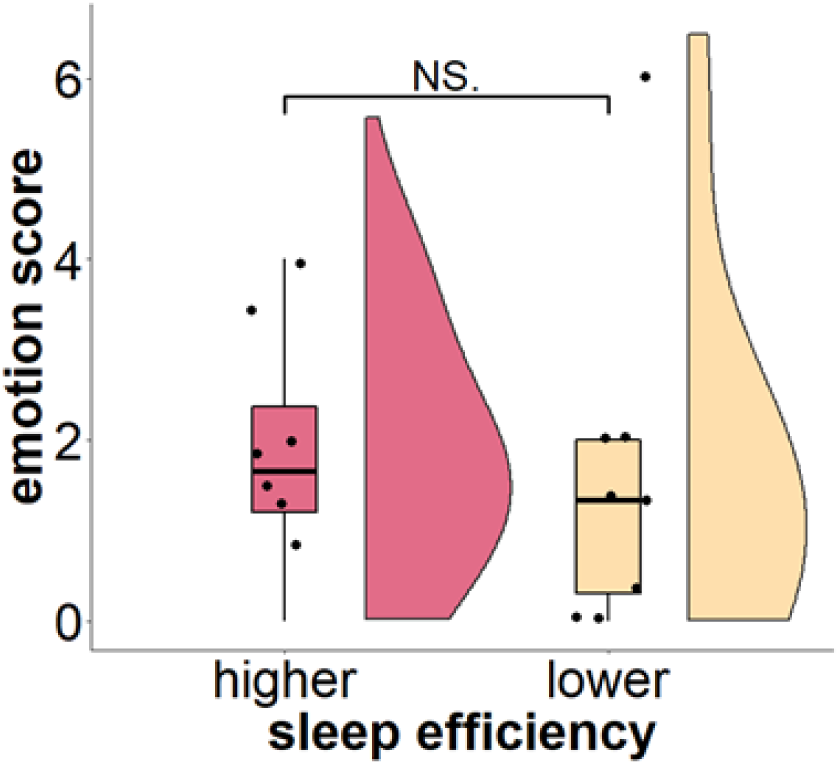
Emotionality in dreams by sleep efficiency. Emotion score in dreams based on participants’ average sleep efficiency across awakenings (Range for higher sleep efficiency group: 79% - 93%; Range for lower sleep efficiency group: 51%- 68%; Mann-Whitney U test: U = 38, P=0.5, CI[-1.19 2.0], N = 16). The difference is not significant (N.S).

## Notes

### Competing Interest Statement

The authors have declared no competing interest.

